# Structure-Property-Processing Correlations of Graphene Bioscaffolds for Proliferation and Differentiation of C2C12 Cells

**DOI:** 10.1101/2023.04.25.538356

**Authors:** Lynn Karriem, Joshua Eixenberger, Stephanie Frahs, Domenica Convertino, Tyler Webb, Twinkle Pandhi, Kari McLaughlin, Ashton Enrriques, Paul Davis, Harish Subbaraman, Camilla Colletti, Julia T. Oxford, David Estrada

## Abstract

Graphene – an atomically thin layer of carbon atoms arranged in a hexagonal lattice – has gained interest as a bioscaffold for tissue engineering due to its exceptional mechanical, electrical, and thermal properties. Graphene’s structure and properties are tightly coupled to synthesis and processing conditions, yet their influence on biomolecular interactions at the graphene-cell interface remains unclear. In this study, C2C12 cells were grown on graphene bioscaffolds with specific structure–property– processing–performance (SP3) correlations. Bioscaffolds were prepared using three different methods - chemical vapor deposition (CVD), sublimation of silicon carbide (SiC), and printing of liquid phase exfoliated graphene. To investigate the biocompatibility of each scaffold, cellular morphology and gene expression patterns were investigated using the bipotential mouse C2C12 cell line. Using a combination of fluorescence microscopy and qRT-PCR, we demonstrate that graphene production methods determine the structural and mechanical properties of the resulting bioscaffold, which in turn determine cell morphology, gene expression patterns, and cell differentiation fate. Therefore, production methods and resultant structure and properties of graphene bioscaffolds must be chosen carefully when considering graphene as a bioscaffold for musculoskeletal tissue engineering.

## Introduction

Musculoskeletal conditions are often debilitating and can cost the US an annual estimated $213 billion in lost wages and medical costs arising from treatments.^1^ These treatments are generally limited to symptomatic relief from invasive surgery or total replacement. Volumetric muscle loss can be especially challenging to treat due, in part, to the absence of the extracellular matrix (ECM), which removes the biophysical and biochemical signaling cues that aid in regeneration.^2^ One potential solution for treating such injuries is through the transplantation of muscle to replace damaged tissue. While this approach has shown some success, transplantation requires a suitable muscle tissue donor or donor site, which may be limited and may create additional injury.^3^ Promising alternatives to transplantation for skeletal muscle regeneration include physical therapy, cell therapy, nanotechnology, and tissue engineering.^4^ While these options may overcome many of the associated challenges, advances are needed to make tissue engineering a truly viable solution.

Tissue engineering approaches involve the utilization of biocompatible scaffolds to support cell growth to replicate the biophysical properties of the native tissue. Bioscaffolds must provide structural support, facilitate cell adhesion, deliver molecular signals such as growth factors, and induce cells to produce ECM that matches the native tissue microenvironment.^2^ Tissue engineering bioscaffolds serve as a template for tissue formation and can be seeded with stem cells and growth factors to augment the healing process.^5^ While there have been tremendous advances in tissue engineering approaches, more research is needed to develop suitable biocompatible scaffolds. In this regard, graphene has emerged as a potential bioscaffold for many tissue engineering applications.

Graphene bioscaffolds have been utilized in conjunction with a variety of cell types, including cardiomyocytes,^6^ neuronal cells,^7, 8^ mesenchymal stem cells,^9^ and myoblasts^10^ to generate numerous different tissues. Graphene bioscaffolds possess many promising attributes including biocompatibility, serum protein adsorption, high tensile strength, and electrical conductivity. Graphene has even been shown to promote differentiation of certain musculoskeletal cell lines.^10-13^

Since the first realization of graphene in 2004, numerous methods to produce graphene have been developed, including micromechanical exfoliation, laser induced graphene, chemical vapor deposition (CVD), liquid phase exfoliation (LPE) of bulk graphite, and epitaxial growth.^14-20^ Each of these methods have their own benefits and limitations, including the resulting quality of graphene, defects, grain size distribution, yield, and the number of layers. True monolayer graphene consists of a single layer of carbon atoms arranged in a hexagonal honeycomb structure. Bilayer or few-layer (3-10 layers) graphene is often used in bioscaffold studies; however, the precise number of graphene layers can radically alter the mechanical, electrical, and thermal properties of the material.^21^

Researchers have explored various geometries of graphene such as patterned islands,^22^ crumpled bioscaffolds,^23^ and CVD hybrids.^24^ Bajaj et al. found that C2C12 cell growth aligned with the graphene patterned islands.^22^ Kim et al. found that their crumpled graphene promoted alignment, elongation, differentiation, and maturation of the C2C12 cells.^23^ Although numerous studies have evaluated graphene bioscaffolds, very few studies focus on the impact of different graphene synthesis methods on the cellular response of specific cell lines, making it difficult to correlate the relationship between graphene analogs and cellular dynamics.

Here we report our evaluations of the structure-properties-processing-performance relationships of graphene bioscaffolds produced using three different synthesis methods, and the resultant effects on the morphology and differentiation of murine C2C12 cells toward myogenesis. A schematic representation of graphene synthesis and our experimental outline is shown in Figure 1.

**Figure 1.**
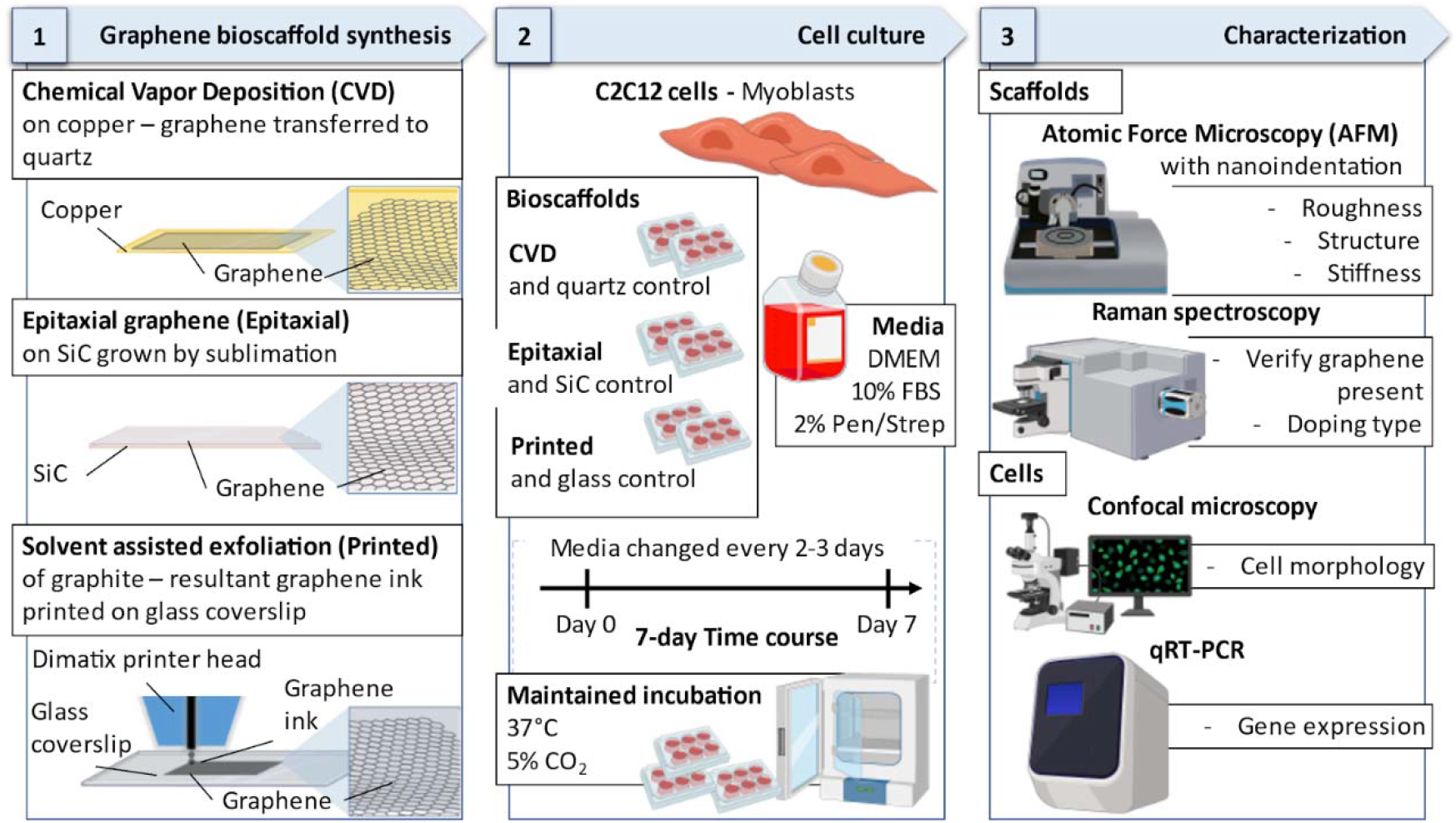
Schematic of the different graphene synthesis methods, cell culture outline, and the characterization techniques utilized.

Graphene bioscaffolds were produced using three different methods - CVD, sublimation of silicon carbide (SiC), and printing of LPE graphene. Each scaffold exhibits distinct physicochemical properties that lead to changes in cellular dynamics and morphologies. Our cell-based studies also reveal distinct gene expression profiles of C2C12 cells grown on each graphene bioscaffold. Collectively, we demonstrate that synthesis methods for the construction of different planar graphene bioscaffolds impact the material’s properties and ultimately cell dynamics.

## Results

### Graphene Characterization

To investigate how variations in graphene’s structure and properties affect cellular growth dynamics, we compared three common methods to produce graphene; graphene grown by CVD on copper foil, epitaxial growth via sublimation on SiC, and LPE. After obtaining graphene, the samples were deposited onto specific substrates or left on the substrate in the case of epitaxial grown graphene. For graphene grown by CVD, the graphene film was removed from the Cu foil by electrochemical delamination and transferred to quartz. Graphene grown via thermal decomposition of SiC was utilized as prepared. Finally, an ink was formulated out of the LPE graphene and deposited on a glass coverslip using inkjet printing.

Atomic force microscopy (AFM) and Raman spectroscopy were employed to characterize the graphene bioscaffolds and identify specific variations in properties that might impact downstream cellular dynamics. Specifically, AFM was utilized to determine the surface morphology, roughness, and elastic modulus of each bioscaffold (Figure 2; see Figures S1-S3 for additional information). Topography maps of three 10 µm x 10 µm areas of each scaffold were acquired and used to calculate the RMS surface roughness (R_q_). CVD and epitaxial graphene (Figure 2a-b) exhibited the most uniform surface, with an associated R_q_ of ∼2 nm. Despite having similar RMS surface roughness, they exhibited extremely different morphologies. SiC graphene (Figure 2b) possessed a layered structure along the surface, with well-defined terraces, whereas the CVD graphene had smaller grains and visible nucleation sites. As expected, the printed graphene (Figure 2c) had the roughest surface, with large peak-to-valley measurements and displayed a RMS surface roughness of 22.6 nm.

**Figure 2.**
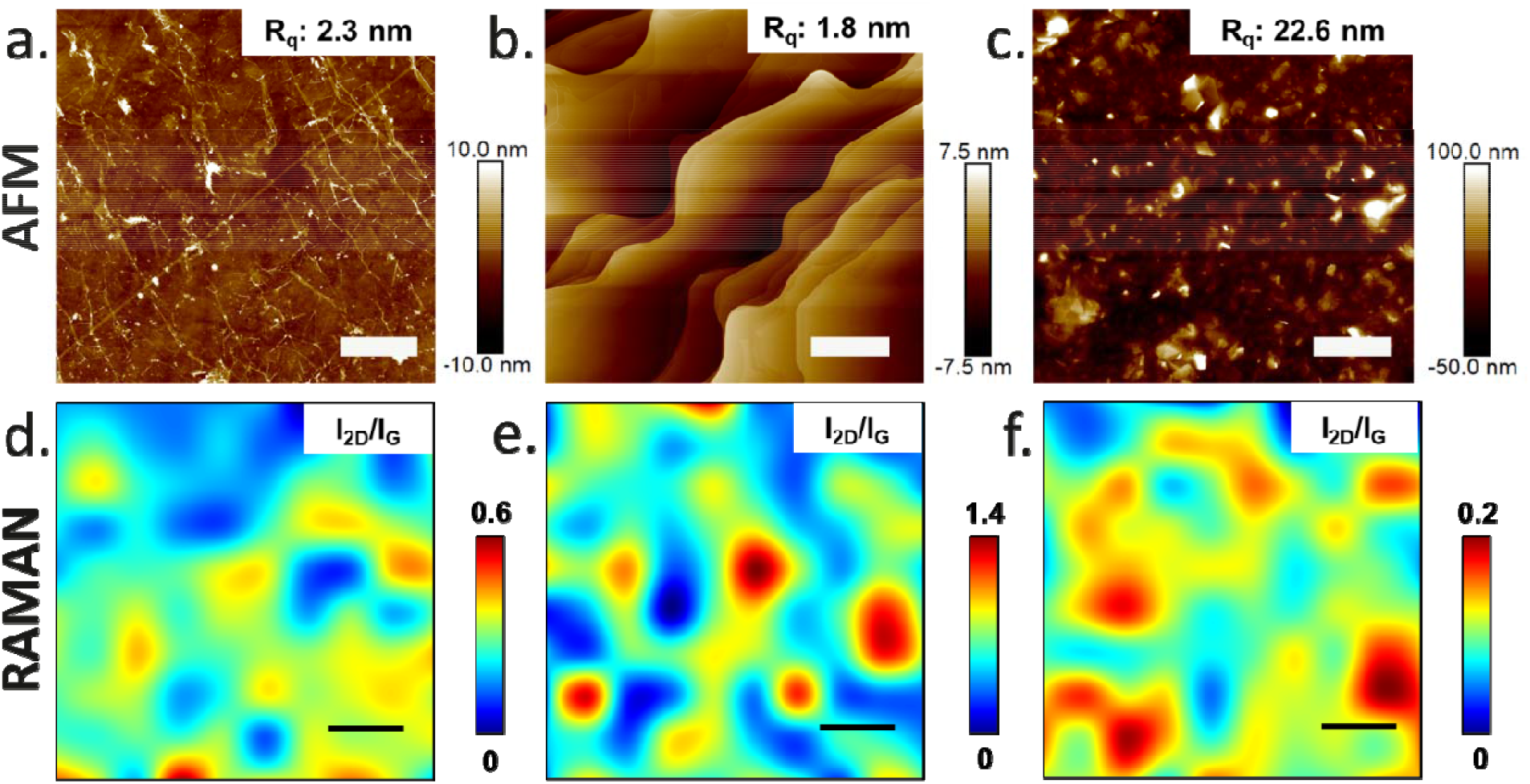
Graphene bioscaffolds characterized by AFM and Raman microspectroscopy. AFM images showed surface structure (morphology) and roughness for a) CVD, b) epitaxial, and c) printed graphene. Raman intensity maps of the 2D to G peak ratios for d) CVD, e) epitaxial, and f) printed graphene. (AFM and Raman map scale bars = 2 µm)

The Raman spectrum of graphene has three characteristic peaks near 1580, 1350, and 2700 cm^-1^ known as the G, D, and 2D peaks, respectively.^25, 26^ The ratio of the intensities of the 2D to G peaks is related to the number of layers in graphene, which can significantly impact its properties.^27, 28^ Raman spectroscopy maps of the 2D (I_2D_) to G (I_G_) peak intensity ratios (I_2D_/I_G_), are shown in Figure 2d-f. The printed graphene sample showed the largest surface features (AFM) along with the lowest 2D to G Raman peak ratio (∼0 - 0.2), indicating multilayered structures throughout the sample. The epitaxial graphene bioscaffold showed the highest I_2D_/I_G_ ratios, ranging from 0 to 1.4, indicating more monolayer to few-layer graphene present compared to the other graphene samples. ^27, 28^ Even though CVD graphene and epitaxial graphene had comparable surface roughness, Raman map ratios indicated that CVD graphene had more layers than epitaxial grown graphene.

The measured range of elastic or Young’s modulus (in GPa) for each bioscaffold is shown in Figure S4. Elastic modulus measurements were obtained by AFM cantilever-based nanoindentation.^29, 30^ Epitaxial graphene (Figure S4) showed the highest average stiffness (69.4 GPa), followed by CVD graphene (12.6 GPa), and printed graphene, which had the lowest value (11.5 GPa) for stiffness. All values obtained are considerably lower than the associated substrates and the theoretical Young’s modulus of graphene, suggesting that the measured values are primarily influenced by the graphene scaffold fabrication method and/or layer number.

### Cell Growth and Morphology

The mouse muscle precursor C2C12 cell line is a well-documented cell line often used to study myoblast proliferation and differentiation, that has been invaluable for *in vitro* studies aiming to understand myogenesis.^31, 32^ To evaluate how the three different graphene bioscaffolds impact cellular growth dynamics, C2C12 cells were seeded on the scaffolds and cultured for seven days. Each tissue culture well containing the graphene scaffold was first coated with agarose gel to promote the cells to only grow on the graphene scaffold so that changes in morphology and gene expression profiles were not impacted by cells growing on tissue culture plastic rather than the scaffold.

Changes in cell morphology in response to the three graphene bioscaffolds are shown in Figure 3. For control experiments, cells were also cultured on the three specific substrates (quartz, SiC, and glass) on which the graphene was deposited. After a seven-day culture, the C2C12 cells displayed dramatically different cell morphologies among the three graphene bioscaffolds and the corresponding controls.

**Figure 3.**
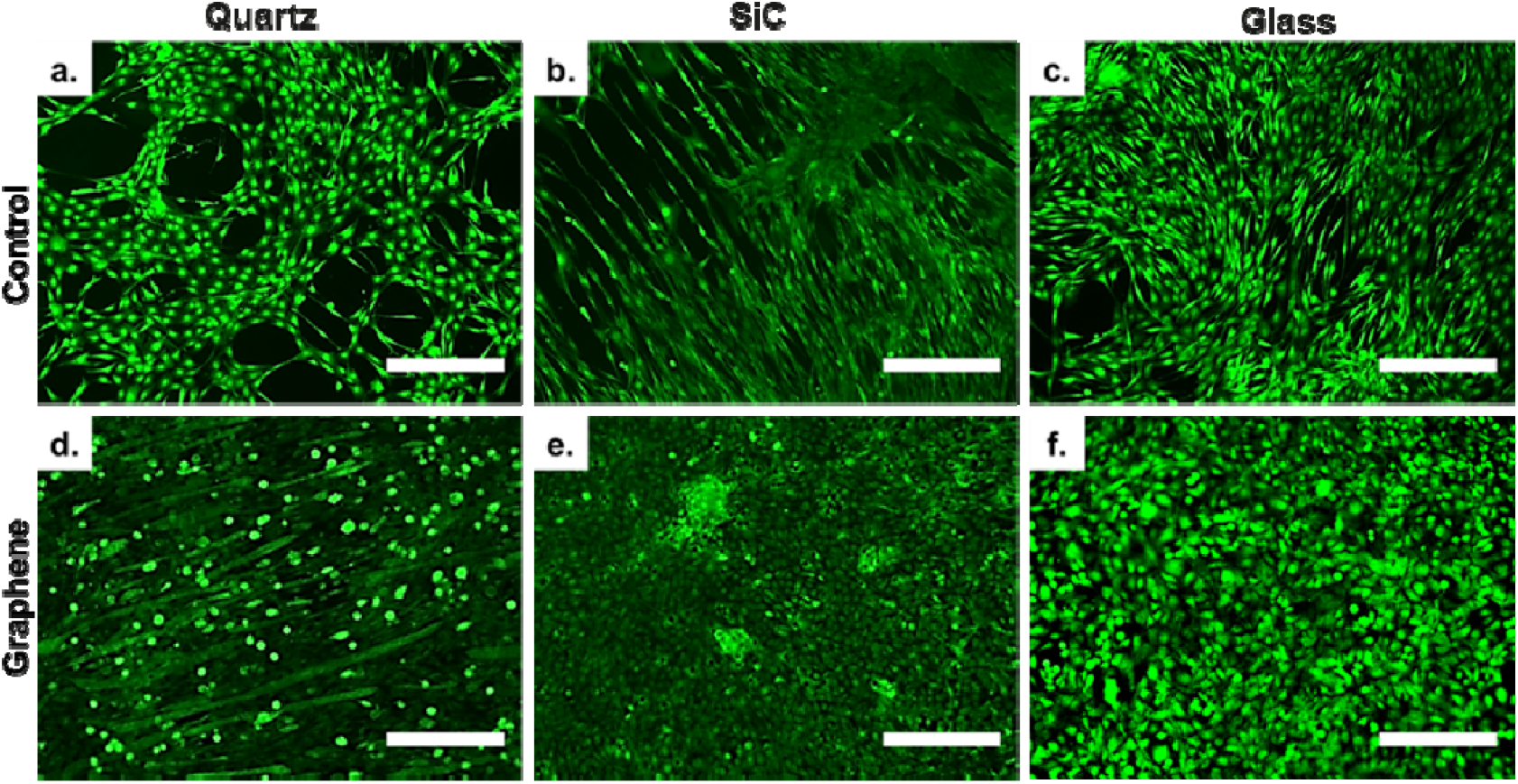
Calcein-AM stained C2C12 images acquired by confocal microscopy show that cell morphology was influenced by graphene bioscaffolds and control substrates. Control substrates: a) Quartz, b) SiC, and c) glass coverslip. Graphene bioscaffolds fabricated by: d) CVD, e) epitaxial, and f) printed LPE graphene. (Scale bar = 300 µm)

The C2C12 cells grown on the control substrates demonstrated substrate specific morphologies. The quartz surface (Figure 3a) resulted in a rounded, fibroblast-like morphology, compared to elongated spindle-like cells on the SiC (Figure 3b) and glass (Figure 3c) substrates. However, distinct morphological changes were observed on the graphene scaffolds when compared to each other and their associated substrate control. The CVD grown graphene bioscaffold on quartz (Fig. 3d) resulted in an elongated cell morphology, with apparent cell fusion, generating multi-nucleate cells in contrast to the corresponding quartz control. Epitaxial graphene bioscaffolds (Fig. 3e) and printed graphene bioscaffolds (Fig. 3f) resulted in cells that had a more rounded, cobblestone-like morphology, which again were distinct from their respective controls. Additionally, epitaxial graphene bioscaffolds supported the formation of tightly packed nodules of cells. No cellular nodules were observed for the cells grown on the printed graphene bioscaffold. The cells grown on the CVD graphene bioscaffold were the only ones to demonstrate an elongated, multinucleate cellular morphology, when compared to the other graphene samples.

Since the cell morphological differences were distinctive and are indicative of cell differentiation lineages, we quantified the differences in morphology by measuring the aspect ratio and alignment of the cells in the different growth conditions. Figure 4 shows the aspect ratios measured for the CVD, epitaxial, and printed graphene bioscaffolds plotted next to their respective control surfaces. Following similar methods as Wang et al., the longest length of the cell structure and the shortest width were measured for 30 cells for each condition (see SI Figure S5-S7).^33^ Aspect ratios for quartz, epitaxial graphene, and printed graphene were all near a value of 1, indicating a spherical shape. The other samples (glass, SiC, and CVD graphene) contained more extended morphologies, all with ratios below 0.2. When comparing the graphene samples, the only one to display a low aspect ratio was the CVD grown bioscaffold.

**Figure 4.**
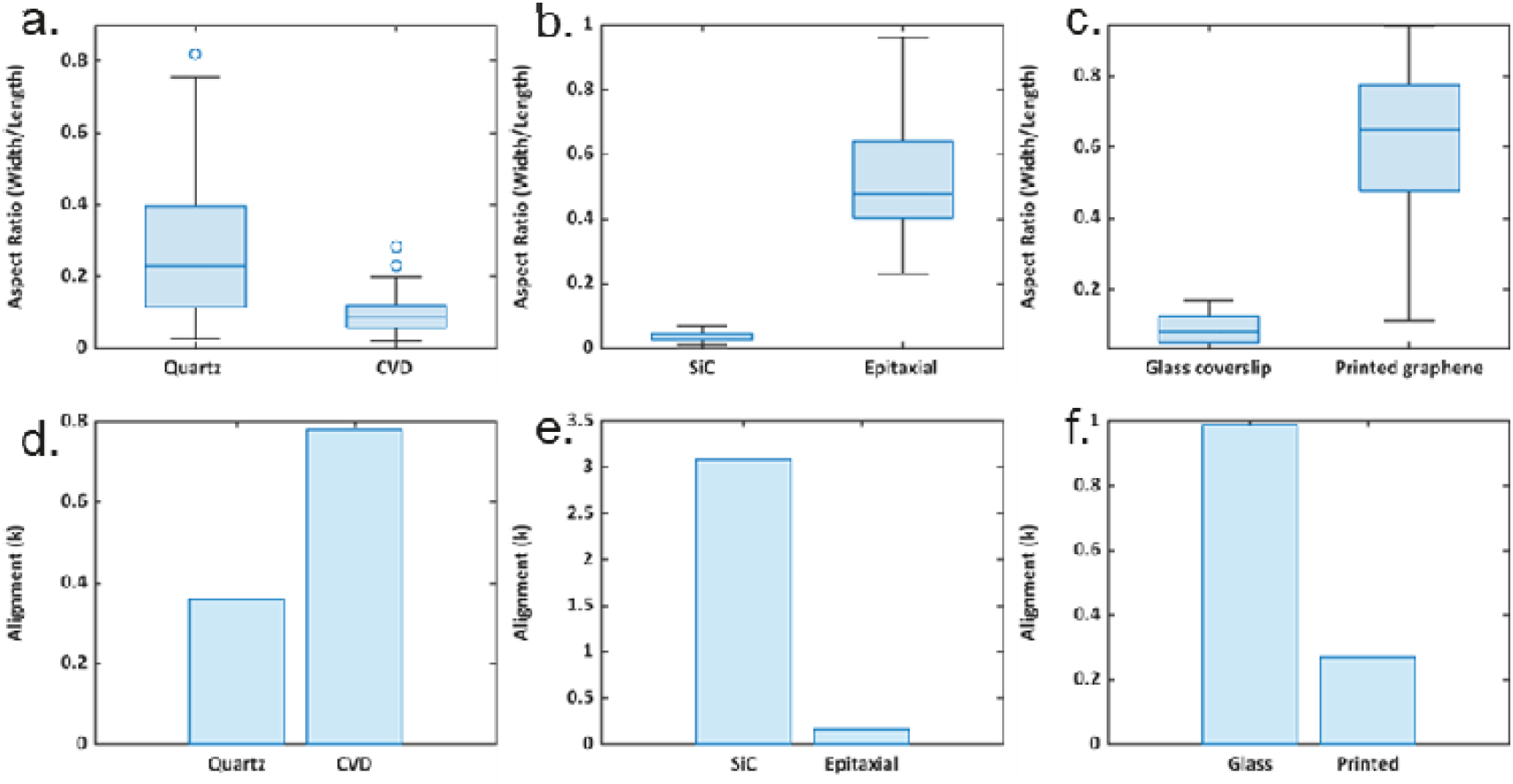
Aspect ratio and alignment studies reveal quantitative morphological differences between the graphene bioscaffolds and respective controls. Aspect ratio for a) Quartz is higher than CVD, b) epitaxial is higher than SiC control, and c) printed is higher than glass. Alignment analysis show increased alignment in d) CVD versus quartz, e) SiC versus epitaxial, and f) glass versus printed.

Muscle cells are typically aligned to form a functioning contractile tissue, and thus cellular alignment is an important factor to consider in tissue engineering applications. To compare the alignment values (k), the CVD, epitaxial, and printed graphene were plotted along with their respective control surfaces (Figure 4d-f). A higher degree of alignment is indicated by larger k-values ^34^. Fiber orientation (µ), sigma, and R^2^ were also calculated (see supporting information S1). All samples showed distinctive differences in their alignment values, with the most significant differences noted between the epitaxial graphene and SiC, which had the highest k-value of all samples measured. Of the three graphene samples, only the CVD graphene had a higher alignment value than its control.

### Gene Expression Profiles

To understand the impact that the graphene scaffold had on the cell’s gene expression profiles, qRT-PCR was performed on the cells grown on graphene scaffolds and compared against the corresponding control substrates. Thirty-five genes related to ECM and muscle cell differentiation were analyzed. Figure 5a shows differential gene expression as a heatmap of log_2_ fold change versus the respective controls. The heatmap includes genes encoding proteins that play a role in attachment, myogenesis, ECM, contractility, and matrix remodeling. For each functional group of genes, a radar plot was created to display gene expression level changes for select genes to compare the differences among the three graphene bioscaffolds (Fig. 5b-f)..

**Figure 5.**
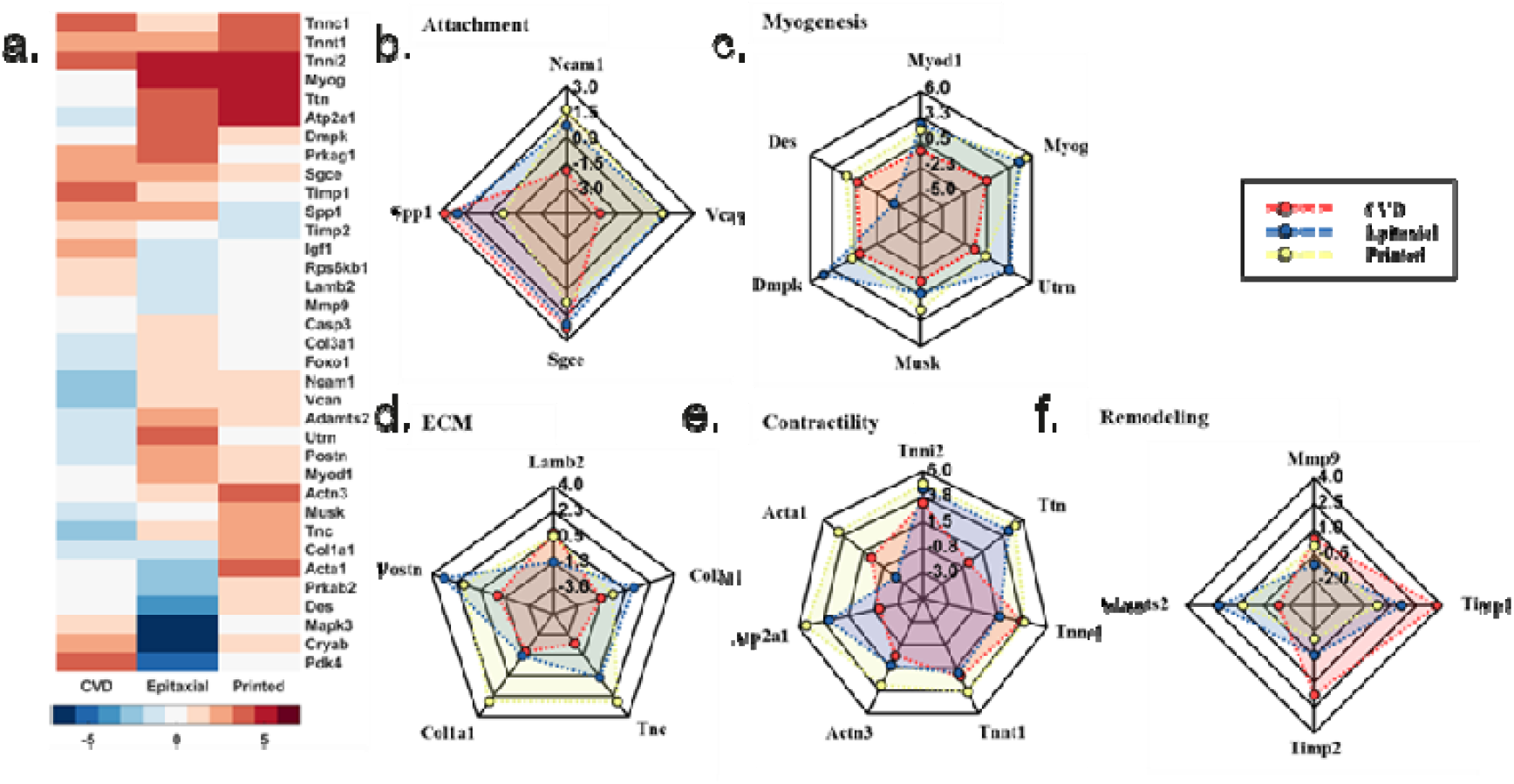
a) Heatmap displaying gene expression fold change (log_2_) for the three graphene bioscaffolds. Radar plots illustrate expression levels of selected genes related to b) attachment, c) myogenesis, d) contractility, e) ECM, and f) matrix remodeling

Since all three graphene samples exhibited different surface roughness, graphene layer numbers, and elastic modulus, it is likely that cellular attachment was impacted. Expression levels of genes related to cell attachment are highlighted in Figure 5b. All three graphene bioscaffolds induced upregulation of *Sgce*, which may indicate that this gene product is related to cellular attachment to graphene in general, regardless of the varying properties of the material. *Ncam1* was downregulated in cells grown on CVD and printed graphene, while *Vcan* was upregulated in cells grown on both epitaxial and printed graphene bioscaffolds. *Spp1* was upregulated in cells grown on CVD and epitaxial graphene, but was downregulated on printed graphene. However, all four genes evaluated were upregulated on the epitaxial graphene bioscaffold.

Six genes involved in myogenesis, *Myod1, Myog, Utrn, Musk, Dmpk*, and *Des* are highlighted in the radar plot (Figure 5c). Each of these genes were upregulated for cells grown on the printed graphene bioscaffold and all but *Des* was upregulated on the epitaxial graphene bioscaffold. Interestingly, even though the morphology of the cells on the CVD graphene appeared the most myotube-like of the three graphene samples, there was only a minor downregulation of two of the genes (*Myog* and *Des*), with the rest mostly unchanged (*Myod1, Utrn, Musk*, and *Dmpk*).

Extracellular matrix (ECM) production from cells is an important factor when designing a bioscaffold to replicate native tissue. To understand the effect of the different scaffolds on ECM production, several genes involved in ECM production were evaluated. *Lamb2, Tnc, Col1a1*, and *Postn* were upregulated on the printed graphene bioscaffold, while *Col3a1* was downregulated. *Col3a1* was upregulated on the epitaxial bioscaffold but downregulated on the CVD grown graphene. *Col3a1* was one of the four out of five genes that was downregulated on the CVD bioscaffold. *Tnc, Col1a1* and *Postn* were also downregulated on CVD graphene, whereas the printed graphene, *Lamb2* was upregulated. *Lamb2* was one of two (*Col1a1*) downregulated genes on the epitaxial bioscaffold – *Col3a1, Tnc*, and *Postn* were upregulated.

Similarly, genes related to matrix remodeling were also highlighted in the radar plots (Figure 5f) *Timp1* was the only gene upregulated in all three bioscaffolds. For CVD and printed bioscaffolds, *Timp1* was upregulated, as was *Mmp9. Timp2* was upregulated on the CVD graphene bioscaffold, but downregulated on the printed graphene bioscaffold. *Adamts5* was upregulated on the printed graphene bioscaffold and downregulated on the CVD bioscaffold. Like the printed bioscaffold, *Timp2* and *Adamts5* were downregulated and upregulated respectively on the epitaxial bioscaffold.

The ability of muscle cells to contract is an essential part of generating forces required for functional muscle tissue. Seven genes related to contractility were evaluated (*Tnni, Ttn, Tnnc1, Tnnt1, Actn3, Atp2a1*, and *Acta1*). All seven of these genes related to contractility were upregulated on the printed graphene bioscaffold, while six out of these seven genes were upregulated on epitaxial graphene (*Acta1 was* downregulated). *Acta1* was upregulated on CVD graphene along with four other contractile genes (Figure 5a). *Atp2a1* and *Ttn* were downregulated on CVD graphene but upregulated on epitaxial and printed graphene bioscaffolds.

## Discussion

Graphene’s use as a bioscaffold for tissue engineering applications has been studied by numerous researchers throughout the field. This versatile material has been shown to be biocompatible with a variety of cell lines and has a wide range of unique properties (e.g., electrical conductivity) not usually present in other polymer-based scaffolds. Interestingly, graphene has even been implicated in inducing differentiation of progenitor cells without traditional growth factors present.^13^

The three graphene synthesis methods resulted in disparate structures and properties. AFM imaging of the surfaces revealed topographical differences, elastic modulus, and RMS surface roughness for each bioscaffold. Raman maps also revealed differences in the quality of graphene when evaluated by the number of layers present. The CVD graphene had a comparable surface roughness to the SiC grown graphene and the lowest standard deviation in the I_2D_/I_G_ peak ratios suggesting a more uniform layer distribution, but the epitaxial graphene had the largest ratio values indicating few layers present. The properties seen in the CVD grown graphene resulted in different cellular morphologies compared to the other two graphene scaffolds. Only the CVD graphene supported elongation and a high degree of cellular alignment, indicating an environment conducive to myotube formation as opposed to the cobblestone-like structures seen on epitaxial and printed graphene. This was further apparent when quantifying the aspect ratio and alignment of the cells. Investigation into gene expression profiles demonstrated that the specific graphene scaffold influences cellular response. Even though some genes related to myogenesis were upregulated in the CVD graphene, the other graphene scaffolds demonstrated upregulation of additional genes related to myogenesis. This could be due, in part, to the fact that the cells grown on the quartz control had aspect ratios and alignment values that were more similar to the graphene scaffold than the other samples evaluated and thus could impact the expression level comparison.

As ECM and muscle related genes were the only genes analyzed in this study, further investigation is warranted to completely understand the impact that the different properties of various graphene scaffolds have on the differentiation of cells. Additionally, other properties of the graphene such as the electronic nature of the material may influence cell growth dynamics. Analysis of the different graphene scaffolds was performed and revealed that the CVD and printed graphene samples appear to be p-type, whereas the epitaxial grown graphene appeared to be n-type (Supporting Info Figure S2),^35^ but more in-depth investigations are needed to understand how this property may impact different cells. While there are other properties and target genes that warrant evaluation, this work demonstrates that variations in graphene properties can dramatically impact cell morphological dynamics and gene expression profiles.

This highlights the need for researchers to fully characterize and report on the graphene properties so that studies aimed at utilizing graphene as a bioscaffold can be more informed and further advance graphene as a viable scaffold for tissue engineering applications.

## Conclusion

This study demonstrated that graphene production and scaffold fabrication methods result in different structures and characteristic properties that ultimately impact cellular morphology and gene expression profiles. Graphene layer number, roughness, elastic modulus, and electronic nature were characterized prior to cell culture. C2C12 murine precursor muscle cells were cultured on three distinct scaffolds, each induced a unique cell morphology and organization after seven days of culture. All three scaffolds caused distinct morphologies and gene expression profiles. The CVD grown graphene appeared to induce a cellular morphology consistent with myotube-like formation, however, this was not necessarily associated with upregulation of the genes involved in myogenesis that were evaluated in this study. Understanding how unique properties of distinct graphene analogs impact differentiation and growth dynamic could prove useful when attempting to produce a specific outcome for tissue engineering applications.

## Materials and Methods

### Chemical vapor deposition (CVD) Graphene film growth and transfer

Graphene was grown on copper (Cu) foil (99.8% Alfa Aesar 0.5 mm) utilizing methane (CH_4_) as a carbon source, similar to previously reported methods.^36^ Briefly, Cu foil was placed in a custom-built 1” tube CVD system.^37^ The system was placed under vacuum and then brought to 1 atm with a continuous ultra-high purity argon (Ar) flow of 1000 sccm while the system heated to 1000 °C. The Cu foil was then annealed in an Ar (800 sccm) and hydrogen (H_2_) (200 sccm) flow for 90 minutes while maintaining 1 atm. To induce graphene growth, the gas switches to CH_4_ (500 sccm) and H_2_ (300 sccm) for 90 minutes at 1000 °C. The resulting graphene/Cu was then cooled to room temperature under an Ar gas flow of 500 sccm and coated with two different layers of poly (methyl methacrylate), or PMMA, to preserve graphene integrity during the transfer process. Both 495 K PMMA A2 and 950 K PMMA A4 were used consecutively to coat the graphene and were baked for 2 minutes at 200°C following each coating. To remove the PMMA/graphene from the Cu foil, the Cu foil was used as a working electrode for electrochemical delamination.^36^ The Cu foil was gradually immersed at a 45° angle in a 0.6 M NaOH electrolyte solution that contained a platinum mesh counter electrode and an Ag/AgCl reference electrode. A -2.1 mV voltage was applied. Once detached, the PMMA/graphene films underwent numerous nanopure water (18.2 MΩ) rinses before transferring to 25.4 × 25.4 mm^2^ quartz glass coverslips. The PMMA was then removed using an acetone bath. Lastly, the resultant graphene/quartz glass composites were annealed at 500°C for 30 minutes under Ar gas flow.

### Graphene Ink Synthesis and Printing

Based on our previously reported methods, we produced graphene ink by solvent assisted exfoliation of bulk graphite powder.^38^ Graphene flakes were obtained by sonicating bulk graphite powder (50 mg/mL) and 2% ethyl cellulose (EC) in ethanol for 90 minutes. The graphene/EC was then centrifuged at 4500 RPM for 60 minutes to remove any remaining graphite and the supernatant was immediately collected. An aqueous solution of NaCl (40 mg/mL; Sigma-Aldrich, >99.5%) was added to the supernatant and centrifuged at 4500 RPM for 15 minutes to collect the graphene. The resulting graphene/EC was then transferred to a PTFE (Teflon) plate and dried overnight. To formulate an ink for inkjet printing (IJP), the dried graphene/EC was then sonicated in a mixture of 92.5% cyclohexanone and 7.5% terpineol solution for 30 minutes. The mixture was then centrifuged at 4500 RPM for 15 minutes to remove any non-dispersed flakes and the supernatant was collected. The resulting graphene ink (3.5 mg/mL) was printed onto glass coverslips using a Dimatix/Fujifilm 2850 IJP system. Graphene was printed using 10 print passes and baked at 350 °C for 20 minutes to remove solvents and residual polymers.

### Epitaxial Graphene Growth on SiC

Epitaxial graphene on SiC was obtained via thermal decomposition of SiC in a commercial cold-wall reactor (BM, Aixtron). Before growth, the 4H-SiC(0001) samples hydrogen etched at 1200-1210 °C and 450 mbar for 5 minutes, to obtain atomically flat surfaces (https://doi.org/10.4028/www.scientific.net/MSF.615-617.589). Monolayer graphene was subsequently grown by heating the samples in Ar atmosphere at 1300 °C and 780 mbar for 5 minutes, similar to what was previously described.^8, 39^ Hydrogen etched SiC(0001) dices were used as control substrates. Sample dimensions were 10 x10 mm^2^.

### Materials Characterization

#### Atomic Force Microscopy

PeakForce tapping mode AFM imaging was performed on the three different graphene bioscaffolds samples using a Bruker Dimension FastScan AFM. Surface topography was mapped using a ScanAsyst-Air-HR probe (Bruker, *k* = 0.4 N/m, 2 nm radius of curvature) and a ∼2 nN PeakForce setpoint. Three 10 µm x 10 µm AFM scan with a resolution 1024 × 1024 and pixel size of ∼10 nm was collected on each sample. Image processing and surface roughness analysis were conducted using NanoScope Analysis Version 1.90 (Bruker). All topographical images were processed with an XY plane fit to remove sample tip and tilt, followed by a first order flatten to remove any line-to-line offsets. The average roughness (Ra) of each image was found using the Nanoscope software. Nanoindentation was used to determine elastic modulus.

#### Raman spectroscopy

A Horiba LabRAM HR Evolution confocal scanning Raman microscope was used to collect Raman spectral maps of the three graphene bioscaffolds. A 633 nm He:Ne laser was used in conjunction with a 100x objective, 1800 gr/mm grating, 70 µm aperture, and 50% neutral density (ND) filter to collect spectra from 1,300-3,000 cm^-1^ on a random 100 µm^2^ area of each bioscaffold. The resulting Raman spectral peaks were fit using LabSpec6 software.

#### Cell Culture

Agarose gel (1%) was applied to the bottom of the 6-well plates and on the border of the three graphene scaffolds to prevent cells from migrating off the bioscaffolds during the experiment. The bioscaffolds were sterilized using UV irradiation for one hour. C2C12 (ATCC) bipotential myoblast cells were cultured in DMEM growth medium containing 10% fetal bovine serum and 100 U/mL penicillin, and 100 μg/mL streptomycin in 5% CO_2_ at 37°C. As controls, C2C12 cells were also cultured on the three different substrates (quartz, SiC, and glass). Each graphene bioscaffold and control was seeded with approximately 13,000 cells/cm^2^. Cells were cultured as indicated above and medium was exchanged every two days. On day 7, the samples were then prepared for imaging or genetic analysis. Viable cells were stained using Calcein-AM and imaged using an EVOS cell imaging system (ThermoFisher Scientific) to investigate morphology. Aspect ratio was determined to quantify cell shape using the methods of Wang et al. by measuring the longest length of the cell structure and the shortest width.^33^ Cell alignment for each bioscaffold was quantified by calculating the k value using FiberFit software (https://www.boisestate.edu/coen-ntm/technology/fiberfit/). ImageJ was used to preprocess images acquired of the C2C12 cells on the six bioscaffolds and converted into 8-bit images for FiberFit preprocessing requirements.^34^ Higher k-values corresponded to higher degree of alignment and as seen in the images; conversely, the less alignment seen, the smaller the k-value.^34^ FiberFit also provided values for fiber orientation (µ), sigma, and R^2^ values.

#### q-RT-PCR

RNA was extracted from each sample following the TRIzol protocol for RNA extraction (Thermo Fisher Scientific). RNA concentration was determined by measuring the absorbance at 260 nm. The RT^2^ First Strand Kit (Qiagen) was used to generate cDNA. Qiagen PCR Arrays were used to quantify gene expression levels of genes related to myogenesis and the extracellular matrix. Genes were amplified by q-RT-PCR using a Roche Lightcycler 96 (Roche).

#### Housekeeping gene (HKG) analysis

The gene expression results were normalized to the average of the two housekeeping genes *actb* (encoding cytoplasmic Actin-1) and *gapdh* (encoding glyceraldehyde 3-phosphate dehydrogenase). These two housekeeping genes were chosen among other candidate housekeeping genes based on constant levels of expression independent of experimental conditions in previously established protocols.^11^ These two genes were selected as the housekeeping genes based on comparison to three other candidates (*b2m, hsp90ab1*, and *gusb*). *Actb* and *gapdh* were found to be stably expressed based on minimal difference of variance (see SI Figure S8). Thirty-one genes were analyzed by calculating the normalized 2^-(ΔCt)^ values to determine relative abundance of gene expression compared to controls, which were C2C12 cells grown on the corresponding base scaffolds, namely silicon carbide, quartz, and glass.

## Author contributions

D.E. conceived of the experiment and supervised the overall execution of the project. L.K. performed cell culture and gene analysis with assistance from S.F. and guidance from J.T.O. L.K, D.C., T.P, T.W., K.P., A.E. performed materials synthesis and characterization under the guidance of H.S., C.C., P.H.D., and D.E. L.K., J.T.O., and J.E. wrote the manuscript. All authors contributed to the figures and data analysis in the manuscript. All authors have read and agreed to the published version of the manuscript.

## Acknowledgements

This work was supported under the National Science Foundation CAREER Award #1848516. The authors acknowledge additional support from the Institutional Development Awards (IDeA) from the National Institute of General Medical Sciences of the National Institutes of Health under Grants #P20GM103408 andP20GM109095. We also acknowledge support from the Biomolecular Research Center, RRID:SCR_019174. J.T.O acknowledges support from the M. J. Murdock Charitable Trust; Lori and Duane Stueckle, and the Idaho State Board of Education. D.E. acknowledges infrastructure support under DE-NE0008677 and joint appointment support under DOE Idaho Operations Office Contract DE-AC07-05ID14517. C.C. acknowledges Progetto THE “Tuscany Health Ecosystem” codice ECS00000017 finanziato dall ’Unione Europea – NextGenerationEU PNRR MUR - M4C2 – Investimento 1.5 - Avviso “Ecosistemi dell ‘Innovazione”.

## Data availability

All data generated or analyzed during this study are included in this published article and its supplementary information files.

