## Supporting Information for "Structure-Property-Processing Correlations of Graphene Bioscaffolds for Proliferation and Differentiation of C2C12 Cells"

**S1 | AFM and Raman Analysis of the 2D FWHM**

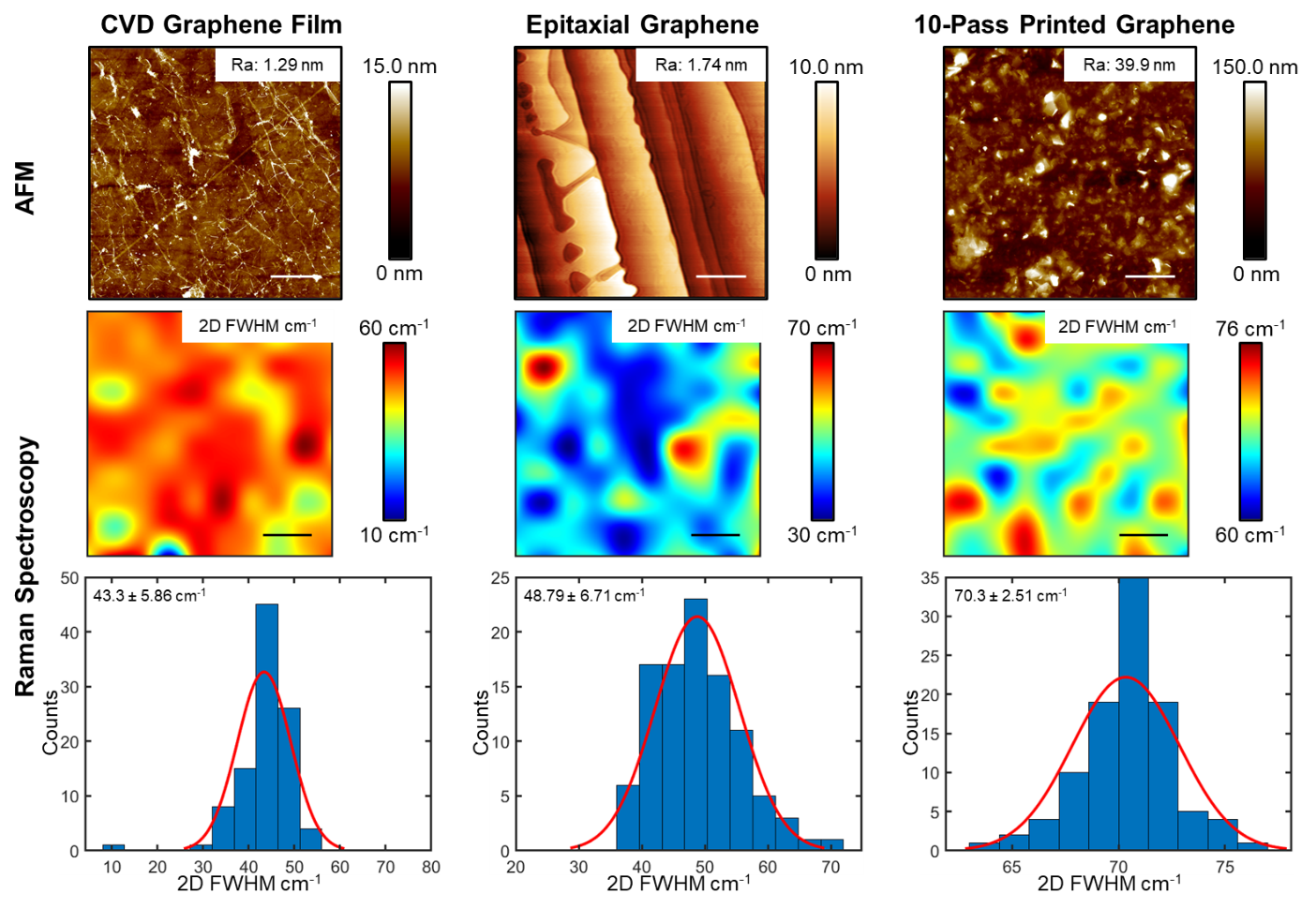

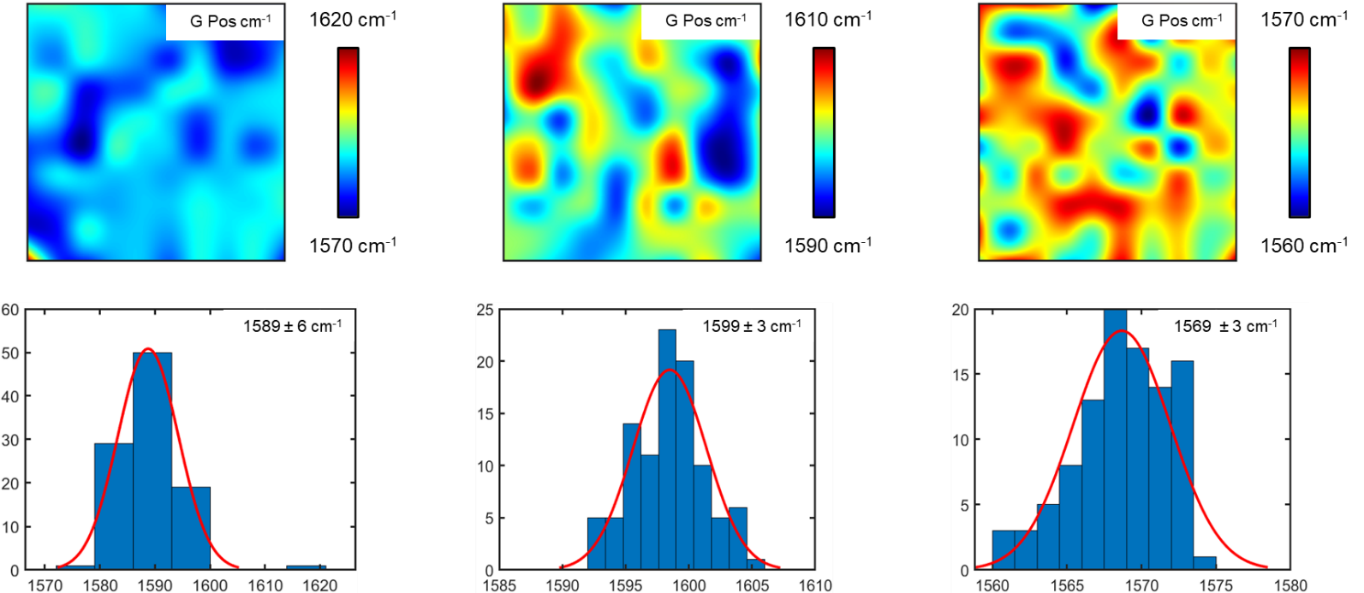

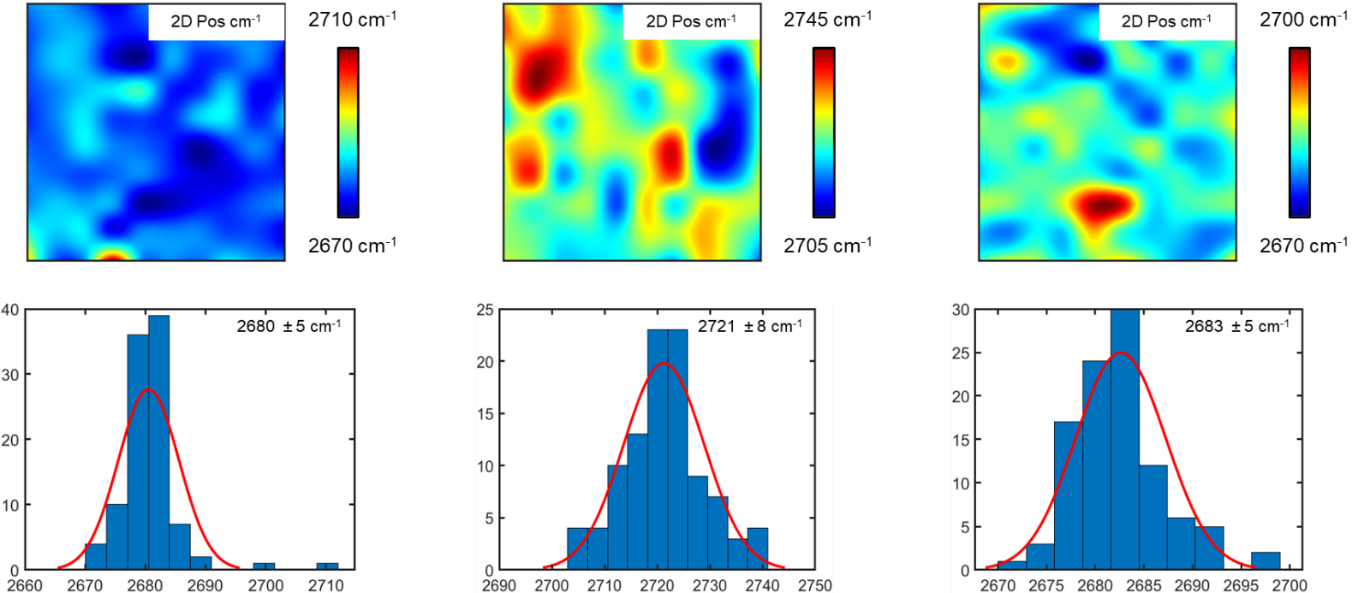
**S2 | Raman maps and histograms of G- and 2D-peak position**

**CVD Graphene**

**10-Pass Printed Graphene**

**Epitaxial Graphene**

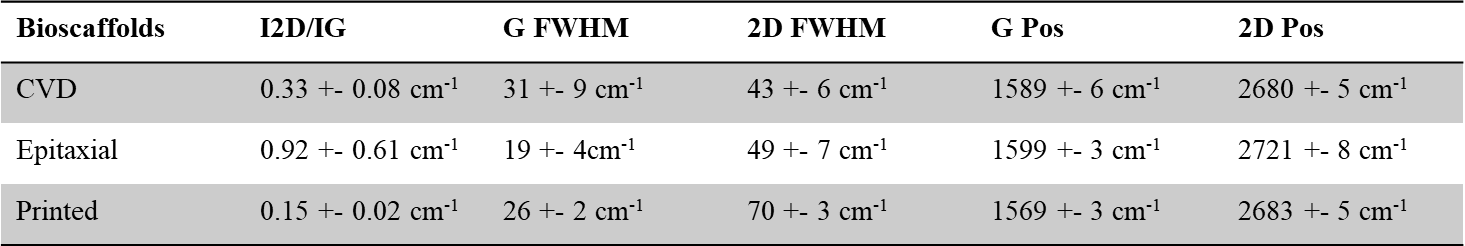
**S3 | Table I2D/IG ratio, FWHM and position for G- and 2D-peaks**

**S4 | AFM Nanoindentation**

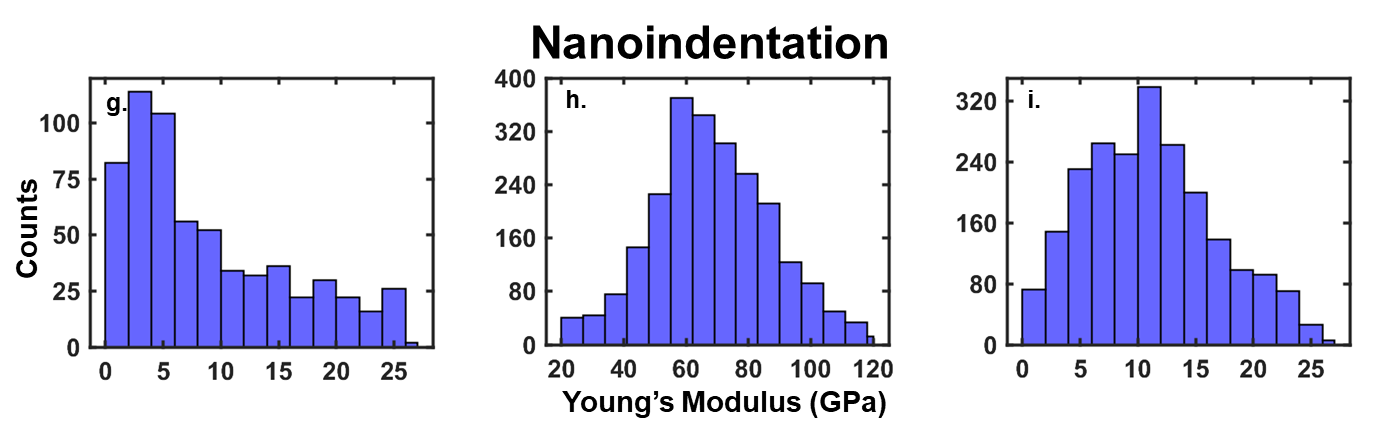

**S5 | Cell measurements**

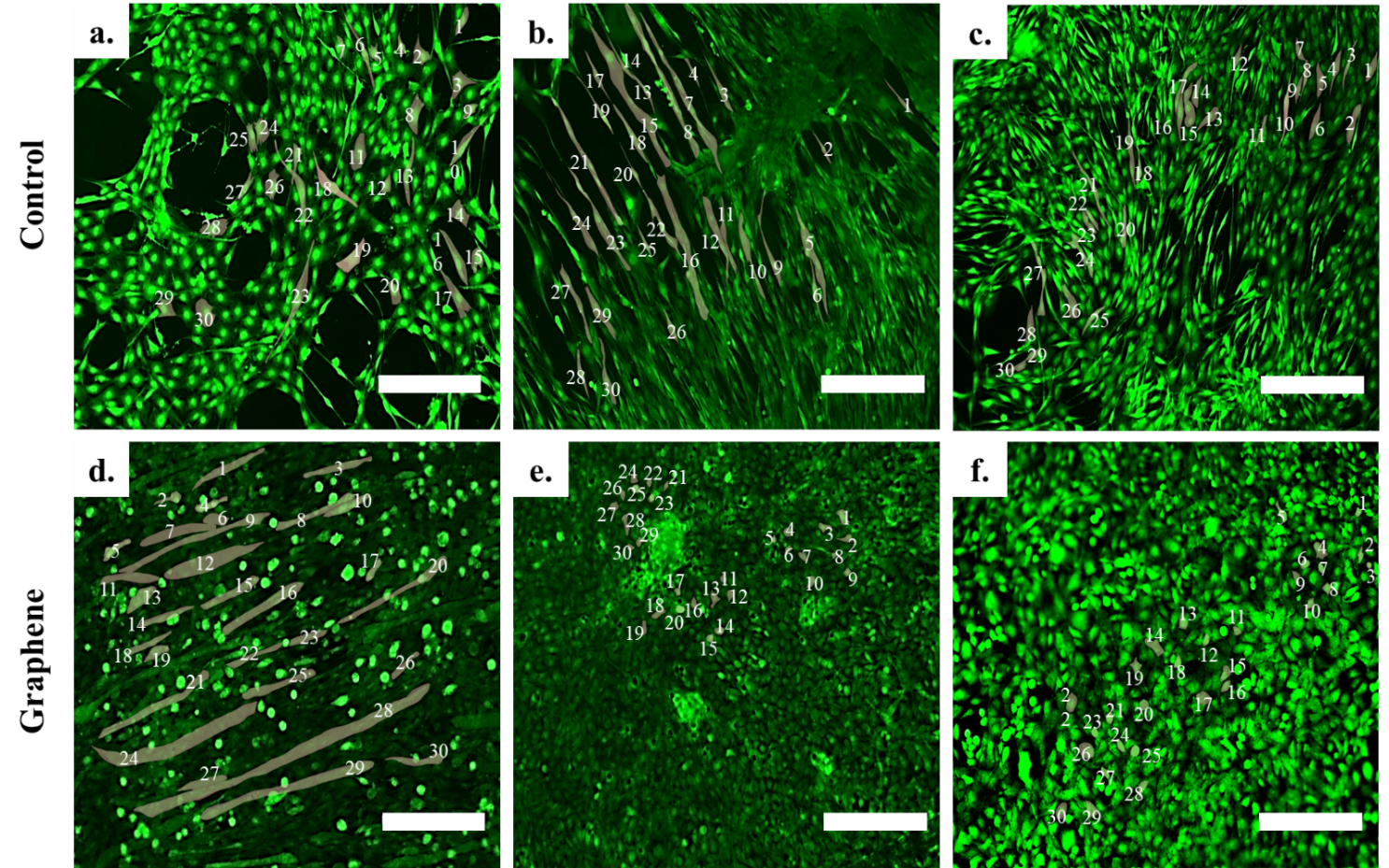

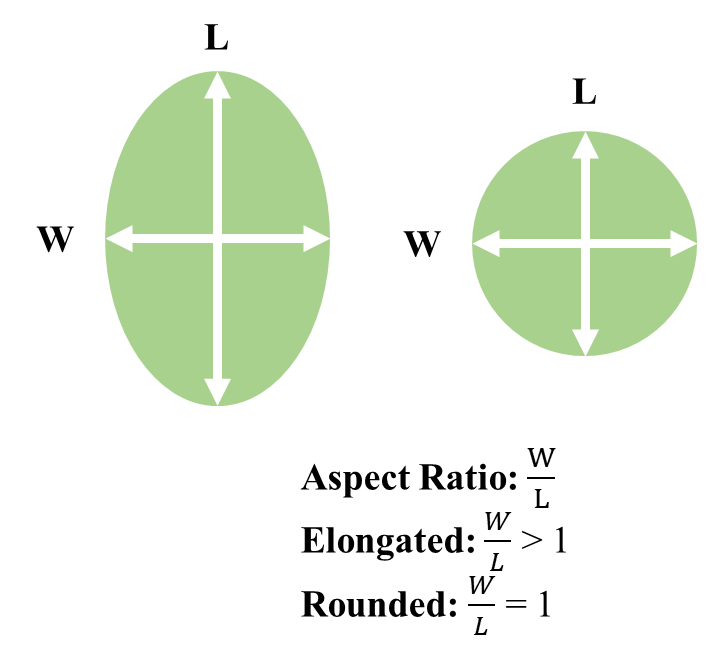

Aspect Ratio: W/L

Elongated: W/L < 1

Rounded: W/L = 1

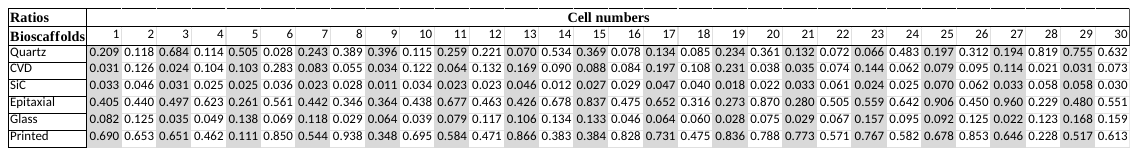

**S6 | Cell aspect ratio**

| Bioscaffold | Mean and STD aspect ratio of cell |
| --- | --- |
| Quartz | 0.2937 ± 0.2214 |
| CVD | 0.0965 ± 0.0616 |
| SiC | 0.0345 ± 0.0152 |
| Epitaxial | 0.5202 ± 0.1932 |
| Glass | 0.0876 ± 0.0438 |
| Printed | 0.6172 ± 0.1990 |

**S7 | Cell alignment**

| Parameter | Quartz | CVD | SiC | Epitaxial | Glass | Printed |
| --- | --- | --- | --- | --- | --- | --- |
| µ (Mean fiber orientation) | 86.06° | 1.23° | -3.78° | 94.76° | 172.11° | 90.24° |
| σ (STD) | 52.84° | 41.32° | 18.1° | 60.01° | 37.03° | 55.81° |
| k (Fiber dispersion parameter) | 0.36 | 0.78 | 3.08 | 0.16 | 0.99 | 0.27 |
| R^2^ | 0.72 | 0.54 | 0.83 | 0.35 | 0.91 | 0.52 |

**S8 | House Keeping Gene (HKG) analysis**

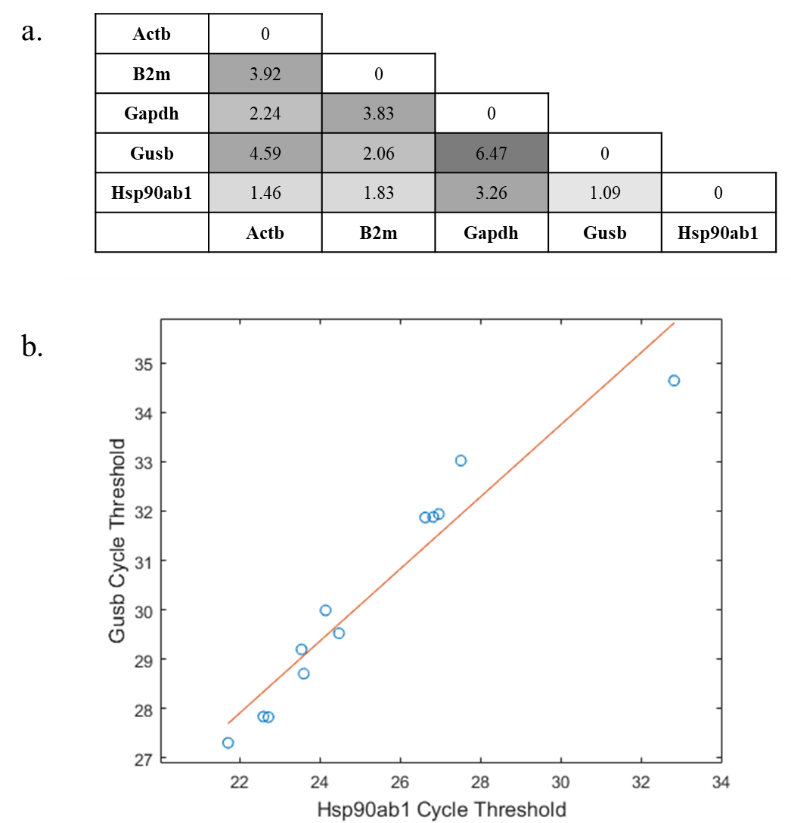

| Graphene Bioscaffolds | | | | | | |
| --- | --- | --- | --- | --- | --- | --- |
|  | Musculoskeletal tissue | Bioscaffold | | Cell line | Gene Expression Analysis (N= No, Y= Yes, I= Intracellular, E=Extracellular) | Reference |
| 1 | Muscle | | Graphene film coated with laminin | Cardiomyocytes | N | (Kim, Kahng et al. 2013) |
| 2 |  | | Graphene/PDMS | C2C12 cells | Yes, I: Green fluorescence protein staining | (Kim, Cho et al. 2016) |
| 3 |  | | Graphene film | C2C12 cells | N | (Bajaj, Rivera et al. 2014) |
| 4 |  | | Crumpled graphene film |  |  | (Kim, Leem et al. 2019) |
| 5 | Bone | | Si/SiO2/Graphene | hMSCs | Y, I: Immunofluorescence staining | (Nayak, Andersen et al. 2011) |
| 6 |  | | Graphene film | C2C12 cells | N | (Crowder, Prasai et al. 2013) |
| 7 |  | | Graphene | (SAOS-2) & hMSCs | Y, I: Immunofluorescence staining | (Kalbacova, Broz et al. 2010) |
| 8 |  | | Graphene film | hASCs & hb=BMMSCs | Y, E: PCR, I: Immunofluorescence staining | (Gu, Lv et al. 2018) |
| 9 |  | | Graphene film & graphene/PDMS | MSCs | Y, E: PCR, I: Immunofluorescence staining | (Xie, Chua et al. 2017) |
| 10 |  | | Graphene film | Dental pulp stem cells (DPSC) | Y, E: PCR, I: Immunofluorescence staining | (Xie, Cao et al. 2019) |
| 11 |  | | Graphene film/PDMS | Human bone marrow derived MSCs | Y, I: Immunofluorescence staining | (Lee, Lim et al. 2011) |
| 12 |  | | Si/SiO2/Graphene | OB-6 |  | (Aryaei, Jayatissa et al. 2014) |
| 13 | Bone-ligament | | Graphene film | PDLSCs & hMSCs | Y, E: PCR, I: Immunofluorescence staining | (Xie, Cao et al. 2015) |
| 14 |  | | Gr/PDMS | hMSC |  | (Ban, Shimoda et al. 2021) |
